# Automated tracking reveals the social network of beach mice and their burrows

**DOI:** 10.1101/2021.08.07.455531

**Authors:** Nicole L. Bedford, Jacob T. Gable, Caroline K. Hu, T. Brock Wooldridge, Nina A. Sokolov, Jean-Marc Lassance, Hopi E. Hoekstra

## Abstract

Evolutionary biologists have long sought to understand the selective pressures driving phenotypic evolution. While most experimental data come from the study of morphological evolution, we know much less about the ultimate drivers of behavioral variation. Among the most striking examples of behavioral evolution are the long, complex burrows constructed by oldfield mice (*Peromyscus polionotus* ssp.). Yet how these mice use burrows in the wild, and whether burrow length may affect fitness, remains unknown. A major barrier to studying behavior in the wild has been the lack of technologies to continuously monitor – in this case, nocturnal and underground – behavior. Here, we designed and implemented a novel radio frequency identification (RFID) system to track patterns of burrow use in a natural population of beach mice. We combine RFID monitoring with burrow measurements, genetic data, and social network analysis to uncover how these monogamous mice use burrows under fully natural ecological and social conditions. We first found that long burrows provide a more stable thermal environment and have higher juvenile activity than short burrows, underscoring the likely importance of long burrows for rearing young. We also find that adult mice consistently use multiple burrows throughout their home range and tend to use the same burrows at the same time as their genetic relatives, suggesting that inclusive fitness benefits may accrue for individuals that construct and maintain multiple burrows. Our study highlights how new automated tracking approaches can provide novel insights into animal behavior in the wild.

## Introduction

Animals construct a diverse array of often stereotyped architectures – ranging from spider webs and fish bowers to bird nests and rodent burrows (Hansell 2005). While some aspects of building behavior may be plastic or learned (e.g., Breen et al. 2016; Hesselberg 2014), it is clear that, in many cases, construction behavior has a genetic component (Dawson et al. 1988; Walsh et al. 2010; Weber et al. 2013; York et al. 2018). In these cases, how and why differences in animal architecture evolve within and among species remains poorly understood. One hypothesis is that variation in animal architecture reflects adaptation to local environment. For example, differences in nest morphology among bird species are often associated with breeding environment (Perez et al. 2020), and variation in web shape among orb-weaving spiders may reflect adaptation to different prey types and habitats (Blackledge and Gillespie 2004).

North American deer mice (genus *Peromyscus*) show considerable natural variation in burrow architecture, and, notably, these interspecific differences have a strong genetic basis (Weber et al. 2013) and correlate with habitat (Hu and Hoekstra 2017; Weber and Hoekstra 2009). For example, oldfield mice (*P. polionotus* ssp.) live exclusively in open habitats and construct long, complex burrows, which may serve fitness-related functions, such as providing refuge from predators (Jackson 2000), places to cache food (Gentry and Smith 1968), buffering against temperature fluctuations (Esher and Wolfe 1979), and/or safe arenas for social interactions (Blanchard et al. 1995). This raises the possibility that burrow evolution in *Peromyscus* mice may be driven by natural selection. However, to determine how burrow architecture may affect survival and/or reproduction within a species, it is first necessary to understand how burrows vary and how they are utilized in the wild.

While burrow building behavior in *Peromyscus* mice has been studied in controlled laboratory settings (Dawson et al. 1988; Metz et al. 2017; Weber and Hoekstra 2009), our understanding of how mice use burrows in the wild remains limited. Early naturalists inferred patterns of burrow use by excavating burrows and counting their occupants (Blair 1951; Rand and Host 1942; Smith 1966), while radiotelemetry has been used to determine burrow location (Van Zant and Wooten 2003). Although these approaches provide snapshots of burrow use, continuous automated recordings in the wild can reveal detailed patterns of burrow use over space and time. Recent advances in animal-tracking technology have enabled activity monitoring with high spatiotemporal resolution and offer new insight into the movement ecology and social dynamics of diverse species (Krause et al. 2013).

Here, we employ a custom radio frequency identification (RFID) tracking system to continuously and non-invasively monitor burrow use in a natural population of Santa Rosa Island beach mice (*P. polionotus leucocephalus*) on the Gulf Coast of Florida. These beach mice are semi-fossorial and dig the longest burrows reported for *Peromyscus* (Hu and Hoekstra 2017), which may be especially important for fitness in exposed beach habitats where vegetation is especially sparse and alternative refuges are scarce (Pries et al. 2009). In this study, we combine RFID monitoring and burrow measurements in the field with genetic analysis in the laboratory to uncover how beach mice use burrows in the wild. By integrating these data, we are able to determine: (1) if there is an association between burrow attributes and mouse activity; (2) the number of burrows a mouse uses, and the number of visitors a burrow has; and (3) the degree of genetic relatedness between mice with similar patterns of burrow use. Together, our findings reveal how beach mice use burrows under fully natural ecological and social conditions and suggest a potential selective mechanism for the evolution of burrow length in *Peromyscus* mice.

## Methods

We monitored burrow use in a single population of Santa Rosa Island beach mice (*P. p. leucocephalus*) on Eglin Airforce Base on the panhandle of Florida over two sampling periods: May 2016 (17 nights) and November 2017 (27 nights). While the results from both sampling periods were consistent (see Supplementary Material), we focus on the findings from our November 2017 field season, which had more timepoints and a larger sample size.

First, to estimate population density and home range size, we conducted a capture-recapture experiment using a grid of 630 Sherman live traps over 16 consecutive nights in November 2017, thus generating a pool of RFID-tagged beach mice (*n* = 32). We collected a small clip of ear tissue for DNA extraction and implanted an RFID tag (Fig. 1A). Next, we released each mouse at its original trap location and noted whether the animal entered a nearby burrow. We flagged these sites, and others we identified while surveying the trap grid, as candidate burrows for subsequent RFID monitoring. Second, we installed custom-built RFID readers at burrow entrances to record the movement of RFID-tagged mice in and out of burrows (Fig. 1B). Over 11 consecutive nights, we recorded RFID activity for 32 different mice at 40 different burrows in the frontal dunes of Santa Rosa Island (Fig. 1C). The locations of successful grid traps and RFID-monitored burrows are shown in Fig. 1D.

**Figure 1:**
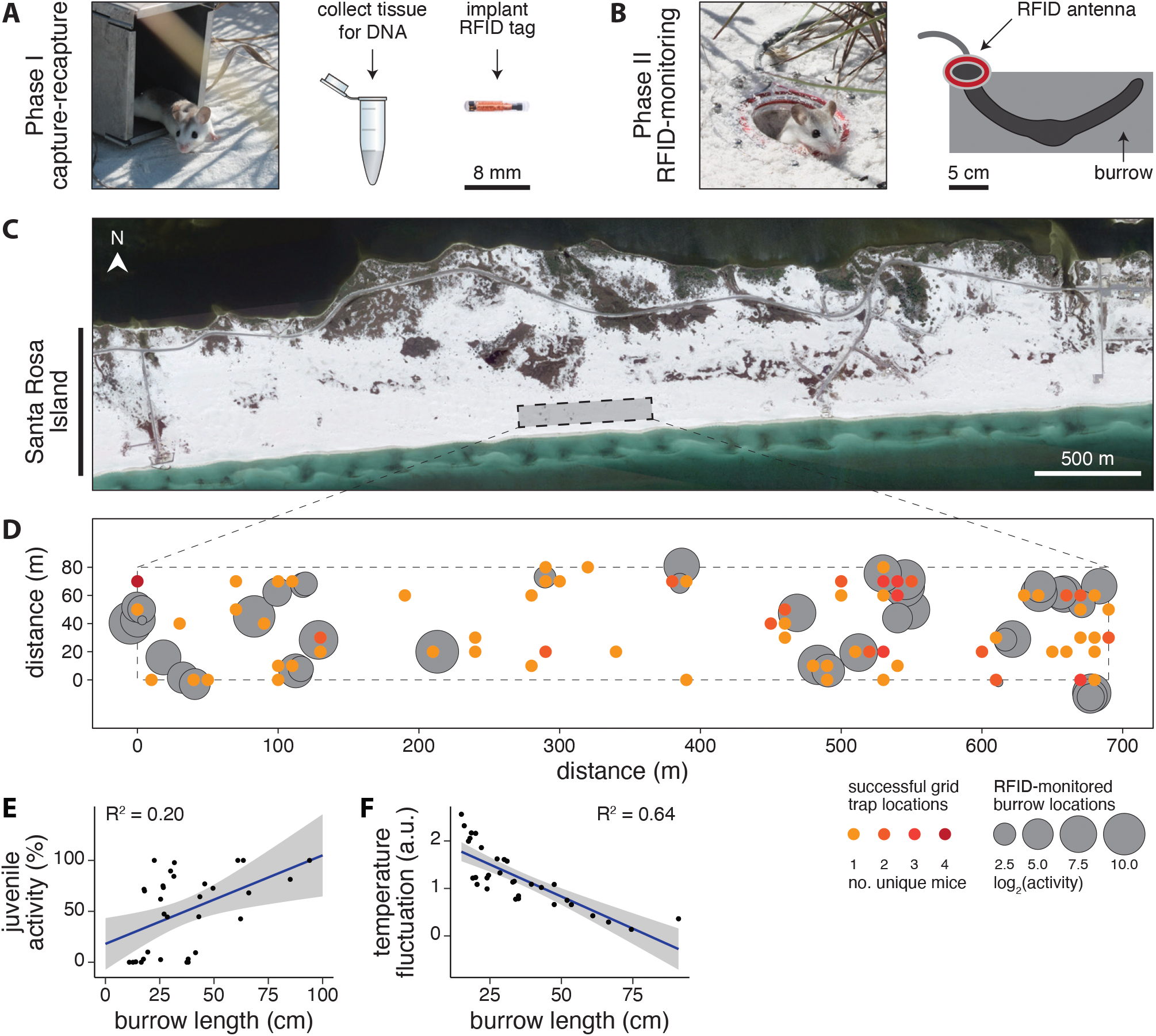
Experimental design and study site. **(A)** *Phase I: capture-recapture*. Photo: beach mouse emerging from a live trap. Prior to release, tissue was collected for DNA extraction and an RFID tag was implanted. **(B)** *Phase II: RFID-monitoring*. Photo: beach mouse exiting a burrow with an RFID antenna installed at the entrance. Typical beach mouse burrow architecture is shown (right). **(C)** Satellite image of Santa Rosa Island, Florida. Dashed line denotes the trap grid boundary. **(D)** Superimposition of Phase I successful grid trap locations and Phase II RFID-monitored burrow locations. Colored circles indicate the number of unique mice caught at each trap location. Grey circles denote the locations of RFID-monitored burrows, log2 scaled by the total number of RFID reads per burrow. **(E)** Relationship between burrow length and the percentage of nightly RFID reads comprised by juvenile mice. **(F)** Relationship between burrow length and temperature fluctuation. For **(E–F)**, data points represent individual burrows, the regression line is shown in blue, and the grey transparency indicates the standard error.

In parallel, we took continuous temperature recordings in natural burrows located outside the trap grid. At the end of the study, we measured the entrance tunnel length of these burrows as well as all RFID-monitored burrows. In addition, we measured the burrow site slope, elevation, heading, and percentage of vegetative cover for all RFID-monitored burrows. In the laboratory, we extracted DNA from ear clips and used a targeted genotyping by sequencing approach to genetically profile all RFID-tagged beach mice. Full experimental details are included in the Supplementary Material.

## Results

### Population sampling

To ascertain the total number of mice at our field site, we fit a spatially explicit capture-recapture (SECR) model to our trap data to estimate population density and home range size (Efford 2018). We derived a population density estimate of 254 mice/km^2^ and a home range size estimate of 14,212 m^2^. These model estimates were consistent with a previous report for this location, which found an average population density of 253 mice/km^2^ and an average home range size of 15,641 m^2^ for beach mice trapped between November 1941 and June 1942 on Santa Rosa Island (Blair 1951). Thus, population density and home range size estimates for Santa Rosa beach mice were remarkably consistent between these two sampling periods separated by more than 75 years.

Next, we used our population density estimate to assess how thoroughly we had sampled the population at our field site. To account for habitat in the vicinity of our traps that might be occupied by beach mice, we included an 80 m habitat mask extending from all edges of our 690 × 80 m trap grid (Fig. S1). In all, we considered 0.204 km^2^ (> 50 acres) of frontal dune habitat in our SECR model. Using our population density estimate, we predicted the total population size at our field site to be 52 individuals. In total, we trapped 43 mice (83% of the estimated total population) and detected RFID activity for 32 of those individuals (62% of the estimated population). Therefore, our RFID data included the majority of predicted individuals at our field site.

### Long burrows have greater thermal buffering and more juvenile activity

After 11 consecutive nights of non-invasive RFID monitoring, we took several burrow site measurements, including the percentage of vegetative cover and the length of the burrow entrance tunnel. Using these data, we tested if any ecological or physical burrow attributes were predictive of nightly mouse activity (i.e., the number of RFID reads per night). First, burrows occurred in areas with no cover to full cover (mean = 58% cover; Fig. S2A), but we found a significant positive relationship between percent cover and nightly activity (R^2^ = 0.14, *P* = 0.010; Fig. S2B), indicating that beach mice primarily use burrows in vegetated patches, while using burrows in open areas less frequently. Next, we tested for an effect of burrow length on nightly activity. While burrows ranged from 11 cm to nearly 100 cm in length (mean = 37 cm; Fig. S2C), we found no association between burrow length and overall nightly activity (*P* = 0.296). However, we did find a higher fraction of juvenile nightly activity at longer burrows (R^2^ = 0.20, *P* = 0.006; Fig. 1E). We also found a significant effect of burrow length on temperature fluctuation, with longer burrows providing greater thermal buffering than shorter ones (R^2^ = 0.64, *P* < 0.001; Fig. 1F, S3). Together, these results indicate that longer, more thermally stable burrows tend to have a higher fraction of juvenile activity than shorter burrows, suggesting they may be natal burrows.

### Mice use multiple, spatially clustered burrows

To understand population-level patterns of burrow use, we performed hierarchical clustering on all 40 burrows, agnostic to spatial location, using mouse visitation data (i.e., the fraction of total RFID activity at each burrow comprised by different mice). This analysis revealed sets of burrows with similar mouse visitation profiles (Fig. 2A). Notably, these burrows tended to be spatially clustered (Komolgrov-Smirnov test, D = 0.96, *P* < 0.001; Fig. 2B, C). Overall, we found that mice used multiple burrows and that burrows were used by multiple mice (Fig. 2D; interactive version available online). Specifically, a single mouse would visit 1-9 burrows per night, and a single burrow would be visited by 1-8 mice per night. Similarly, over the 11-night sampling period, individual mice visited 1-9 burrows (median = 5) and individual burrows were visited by 1-10 mice (median = 3; Fig. 2E, F). We found that mice, on average, revisited 63% of the burrows in their network on a subsequent night. We next asked if mice visited all burrows in their network with equal frequency. Notably, for a given mouse, total RFID activity was not evenly split among burrows. On average, 65% of an individual’s total RFID activity was observed at a primary burrow, while an additional 15% was observed at a secondary burrow (Fig. 2G). The remaining activity was partitioned among 3-9 additional burrows. Therefore, while most of an individual’s nighttime activity was often observed at a single burrow, we found that beach mice consistently use a set of multiple, spatially clustered burrows, both within and across nights.

**Figure 2:**
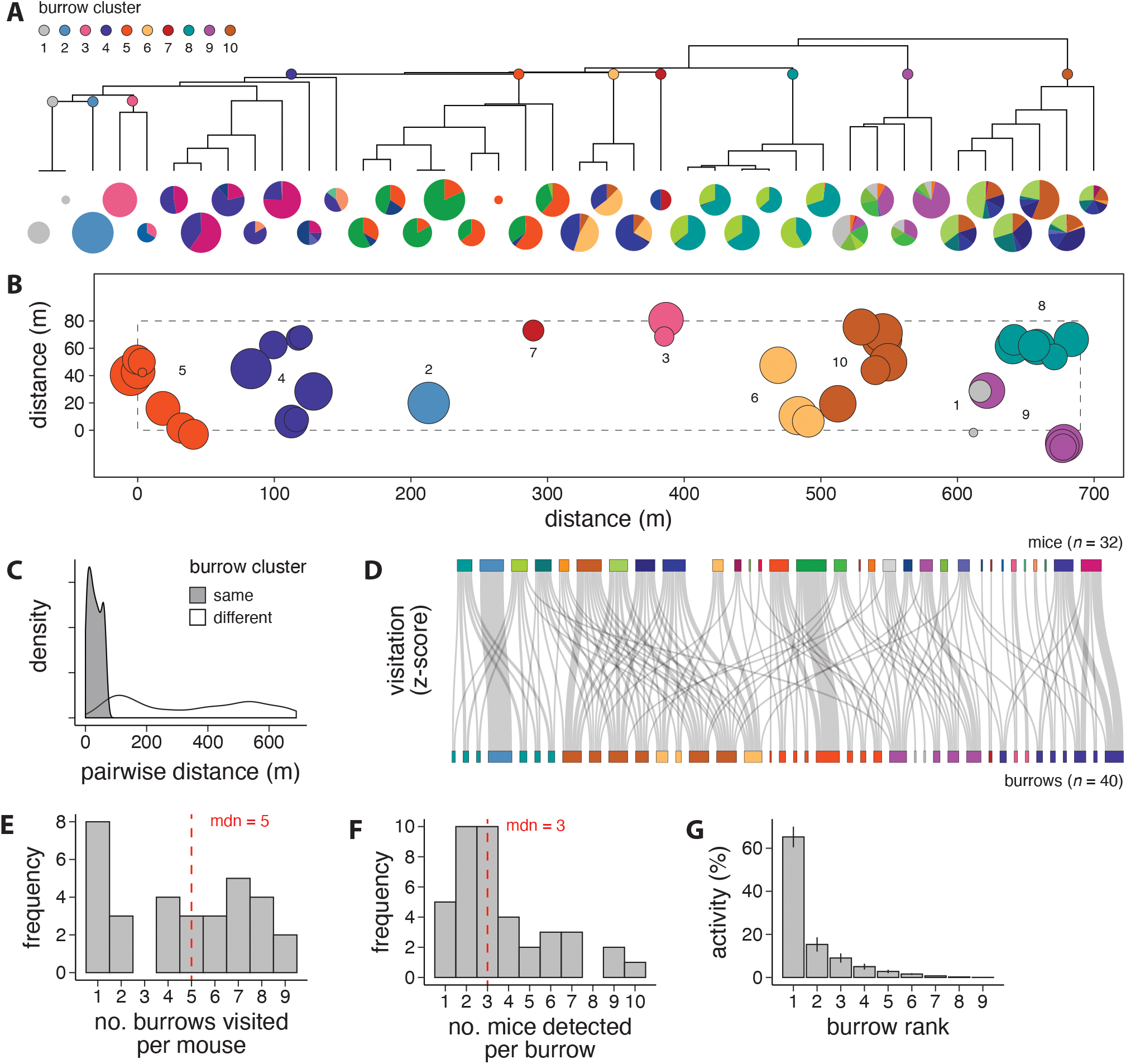
Groups of mice use multiple, spatially clustered burrows. **(A)** Hierarchical clustering of burrows according to mouse visitation data. Pie charts represent individual burrows (*n* = 40), and pie slices represent the fraction of total RFID activity at each burrow comprised by different mice (*n* = 32) over the 11-night sampling period. **(B)** Spatial locations of burrow clusters identified in **(A)**. In **(A)** and **(B)**, circle size is log2 scaled by the total number of RFID reads per burrow. **(C)** Distribtuion of pairwise distances between burrows belonging to the same *versus* different clusters. **(D)** Sankey diagram of mouse visitation data. Individual mice (top) and burrows (bottom) are color-coded as in panels A and B. Grey band widths are proportional to the total number of RFID reads per mouse, per burrow (z-scored). **(E)** Total number of burrows visited per mouse (*n* = 32 mice), and **(F)** total number of mice detected per burrow (*n* = 40 burrows) over the 11-night sampling period. **(G)** Distribution of total RFID activity per mouse, sorted by burrow rank. Error bars represent standard error of the mean.

### Mice show stronger spatiotemporal overlap than expected by chance

We next investigated temporal patterns of burrow use. *Peromyscus* mice have been considered crepuscular, with peak activity at dusk and dawn (Falls 1968). In contrast, we detected ample RFID activity at burrow entrances throughout the night, not limited to dusk and dawn (Fig. S4A). We did, however, find that mice emerged from their daytime burrow (i.e., the burrow where the mouse slept) an hour after sunset and returned an hour before sunrise, on average (Fig. S4B). In addition, mice made several forays throughout the night, both back to their daytime burrow and to other burrows in their network (Fig. S4C).

Given that groups of mice use similar burrow networks, we next assessed whether mice visit the same burrows within the same time windows. Using 90-min sliding window time bins, we calculated an overlap index (OI) for every possible pair of mice and found spatiotemporal overlap (i.e., non-zero OI values) for 74/496 possible pairs (15%). We then performed hierarchical clustering on the matrix of pairwise OI values to visualize groups of mice with similar spatiotemporal patterns (Fig. 3A). While many individuals associated only weakly with other mice in the population, we identified three groups of two, one group of four, and one group of five mice that showed strong spatiotemporal associations (white boxes, Fig. 3A). These groups of mice may represent social units that live together in the same burrow, use the same temporary refuges during foraging forays, or both.

**Figure 3:**
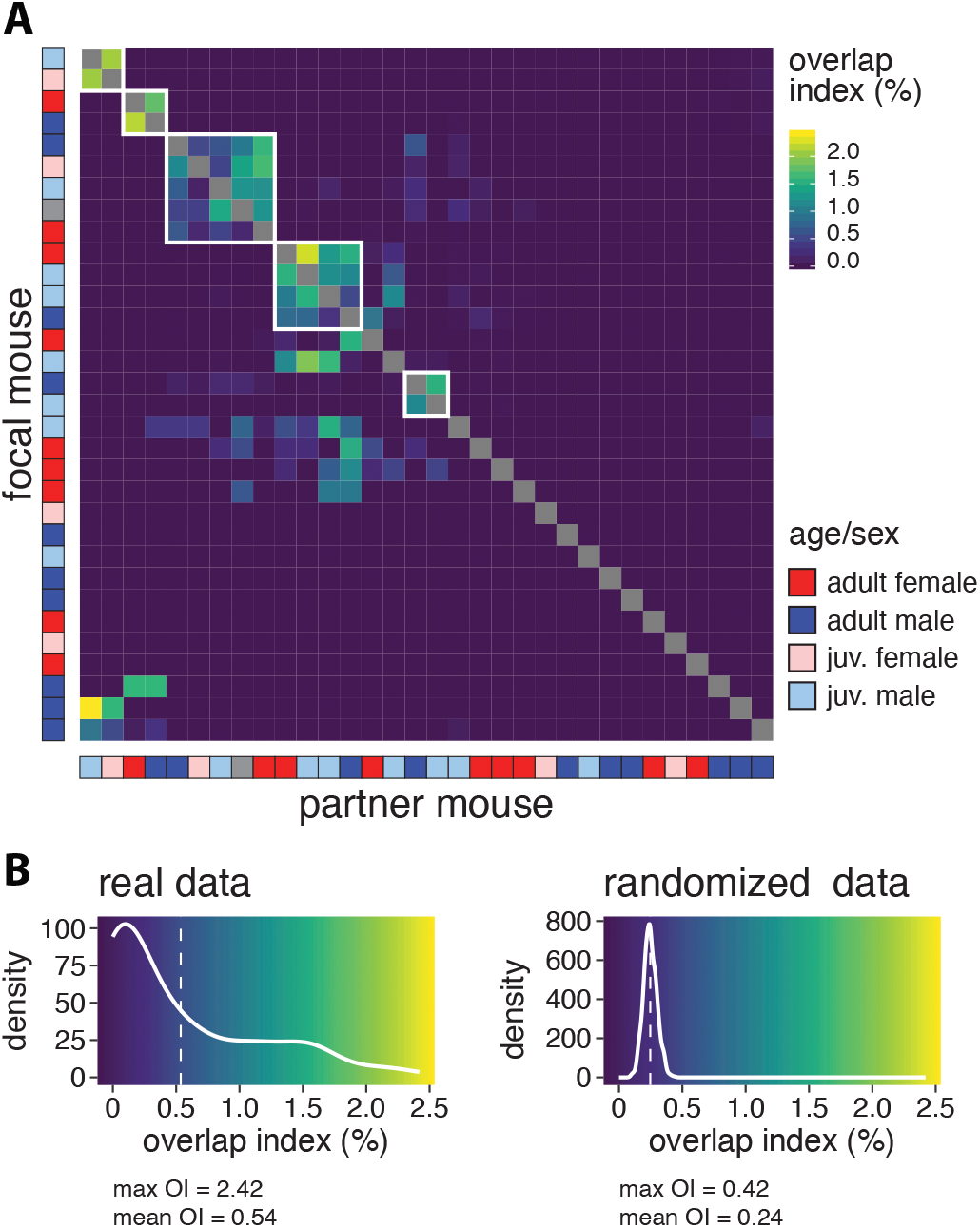
Mice show stronger spatiotemporal overlap than expected by chance. **(A)** Overlap index (OI) matrix for all possible focal and partner mouse pairs. For each focal mouse, OI indicates the percentage of total RFID activity that occurs at the same burrow and within the same 90-min time bin as the partner mouse. **(B)** Distributions of non-zero OIs from real data (left) and randomized data (right). Dashed white lines indicate distribution means.

To test if mice showed stronger spatiotemporal overlap than expected by chance, we randomly assigned mouse and burrow labels to each RFID timestamp and re-calculated OI. When we randomized the data, we found non-zero OI values for 496/496 pairs (100%). However, the distributions of OI values based on real and randomized data differed significantly (Komolgrov-Smirnov test, D = 0.44, *P* < 0.001; Fig. 3B). Specifically, the mean and maximum OI values were considerably larger for real compared to randomized data (Fig. 3B). These findings suggest that, although fewer pairs show overlap than expected from a random model of mouse movement, when mice do overlap, they show stronger spatiotemporal associations than expected by chance. Therefore, we find that mice do not move through the landscape independently, but rather tend to use the same burrows, within the same time frame, as select other mice.

### Genetic relatedness predicts spatiotemporal overlap

We next determined if groups of mice with similar patterns of burrow use correspond to family units. To that end, we generated genetic data at 110 single nucleotide polymorphism (SNP) markers (see Supplementary Material) and estimated relatedness among individuals. First, we visualized a network graph based on genetic relatedness and calculated degree centrality (i.e., the number of direct connections) for each node (i.e., mouse) in the network (Fig. 4A). On average, mice were genetically related to 18 other individuals in the RFID-tagged population, with relatedness coefficients ranging from 0 to 0.75 (mean = 0.15 ± 0.008). We found no significant difference in degree centrality between juveniles and adults or between males and females (age: *P* = 0.142, sex: *P* = 0.325). These findings are consistent with the observation that beach mice settle within a few hundred meters of their natal sites (Swilling Jr and Wooten 2002) and with the prediction that monogamous mammals (like *P. polionotus* ssp.) do not have sex-biased dispersal (Dobson 1982). Second, we visualized a network graph based on OI strength and found that mice, on average, showed spatiotemporal overlap with four other individuals (Fig. 4B). Again, we found no significant difference in degree centrality between juveniles and adults or between males and females (age: *P* = 0.472, sex: *P* = 0.309). Finally, among the pairs of mice that showed spatiotemporal overlap (*n* = 74 pairs), we found a significant positive relationship between genetic relatedness and OI (R^2^ = 0.23, *P* = 0.003; Fig. 4C).

**Figure 4:**
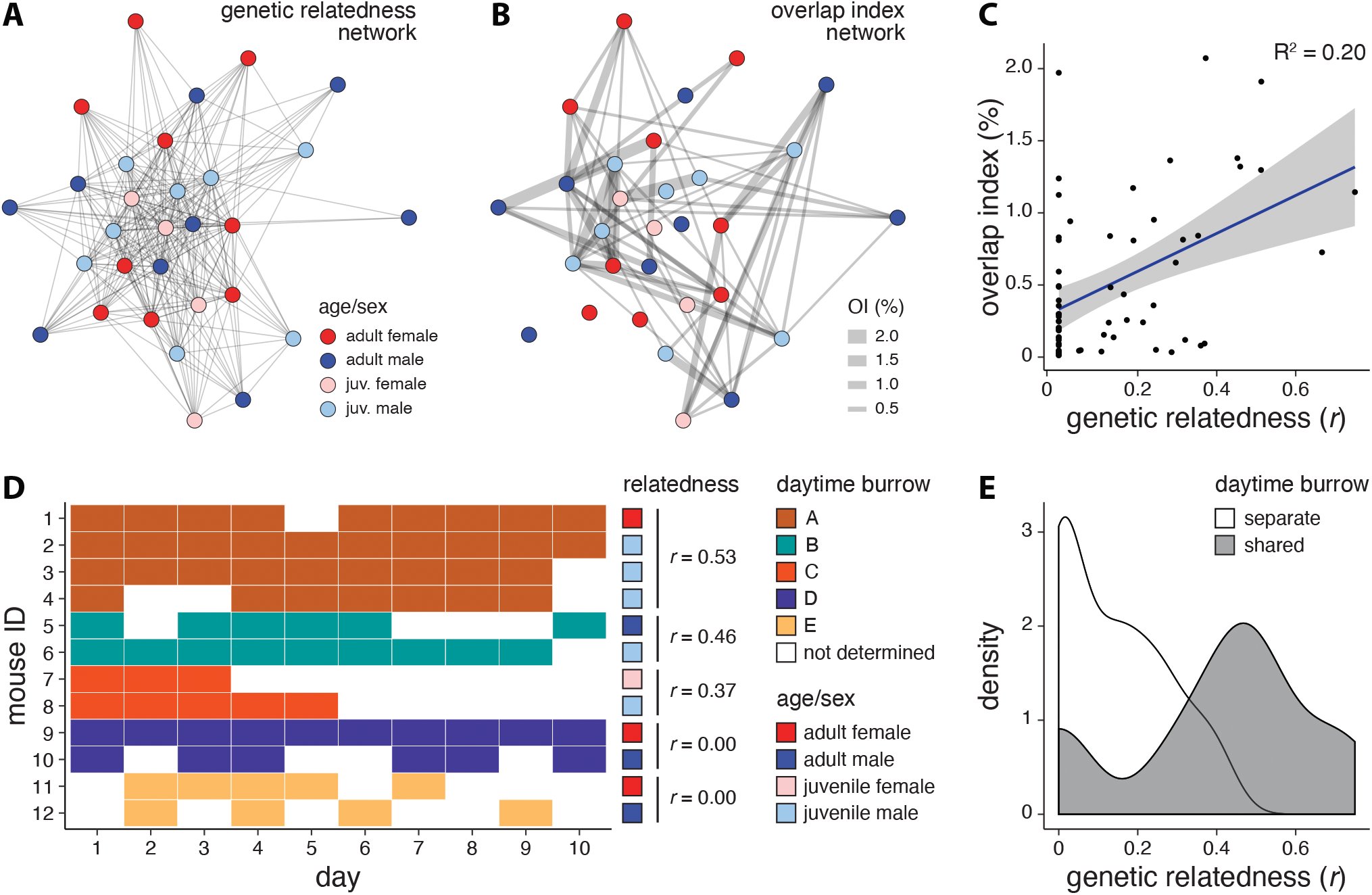
Genetic relatedness predicts spatiotemporal overlap between mice. **(A)** Network diagram of genetic relatedness between mice, with nodes (i.e. mice) positioned by a force-directed algorithm. **(B)** Network diagram of mean overlap index (OI) values. Nodes are positioned as in **(A)** to facilitate comparison, and edges are weighted by OI strength. **(C)** Relationship between genetic relatedness and mean OI for mouse pairs with non-zero OIs (*n* = 74 pairs). Regression line is shown in blue, and grey transparency indicates the standard error. **(D)** Daytime burrow occupancy. Mice 1-12 and burrows A-E were active on all 11 nights. The mean genetic relatedness (*r*) of mice sharing a daytime burrow is indicated (right). **(E)** Distribution of genetic relatedness values (*r*) for mice occupying separate versus shared daytime burrows.

Given that mice were more likely to use the same burrow at the same time as kin, we next ascertained whether related individuals also slept in the same burrow during the day. We restricted our analysis to a subset of 12 mice that were detected on all 11 nights. We found that mice repeatedly returned to the same burrow to sleep – although not on all days, indicating that some mice used at least two different daytime burrows over the 11-night sampling period (Fig. 4D). Nevertheless, apart from two unrelated male-female pairs, individuals that slept in the same burrow during the day had significantly higher genetic relatedness than mice occupying separate daytime burrows (Komolgrov-Smirnov test, D = 0.63, *P* = 0.002; Fig. 4E). Moreover, these daytime burrows had higher percent cover (*P* = 0.012) and higher nightly mouse activity (*P* < 0.001), but the same number of unique mouse visitors (*P* = 0.299) as other RFID-monitored burrows (Wilcoxon rank sum tests). Altogether, these findings suggest that groups of genetically related mice often sleep in the same burrow during the day and use overlapping burrow networks at night.

## Discussion

RFID technology is increasingly used to study rodent social behavior in controlled laboratory settings (Freund et al. 2013; Howerton et al. 2012; Peleh et al. 2019; Weissbrod et al. 2013) and semi-natural to natural environments (König et al. 2015; Smith et al. 2018). However, many of these studies employ artificial habitats, burrows or nest boxes, while fewer record RFID activity at natural features of the environment. To better understand the selective pressures underlying the evolution of burrows and their use, we quantified burrow visitation patterns in wild Santa Rosa beach mice using RFID readers installed at the entrances of their natural burrows. With these data, we made several key findings about the otherwise largely unobservable (i.e., nocturnal and underground) social behavior of these monogamous mice.

First, although we found no association between burrow length and overall RFID activity, we did record more juvenile RFID activity at longer burrows than at shorter ones, suggesting these long burrows may be natal burrows. Notably, we also showed that longer burrows provide a more stable thermal environment, which may be especially important for pup survival, particularly in open habitat with little vegetative cover to dampen thermal fluctuations. At birth, pups have little capacity for independent thermoregulation (Berry and Bronson 1992) and rely on heat retained by the nest for survival (Berry 1970; Brown 1953). Thus, the advantage of a long burrow with greater thermal buffering may be particularly acute at early life history stages, favoring the construction of long burrows by both parents, who care for their young in this monogamous species (Bendesky et al. 2017). Indeed, we have shown in the laboratory that, in *P. polionotus*, opposite-sex pairs with reproductive potential cooperatively build longer burrows than individuals or same-sex pairs (Bedford et al. 2019). Thus, it is possible that natural selection favors a stable thermal environment, particularly in the context of rearing young (i.e., reproductive success), thereby acting as a driver of the evolution of long burrows in this species.

Second, our continuous RFID recordings show that mice visit multiple burrows – up to nine different burrows, both per night and across nights. Moreover, this value likely represents an underestimate, given that we were unable to place an RFID reader on every burrow at our field site. This finding contrasts with previous radiotelemetry data suggesting that beach mice use only one or two burrows (Van Zant and Wooten 2003). However, our data also show that not all burrows are used equally, and that a majority of mouse activity is concentrated at only one or two burrows. In addition, we found a positive relationship between burrow-site percent cover and mouse activity, suggesting that mice may build primary burrows in areas with more vegetative cover, possibly because plant cover aids in predator evasion (Kotler 1984) or because plant root systems stabilize dunes (Stallins and Parker 2003), thereby reducing the probability of burrow collapse. Moreover, the observation that mice rarely visit burrows with low plant cover raises the possibility that mice use burrows in open areas only as temporary refuges during foraging bouts as they move between vegetated patches (Pries et al. 2009; Wilkinson et al. 2013). These results suggest that mice build and use a network of several burrows within their home range, and that these burrows may serve different functions, depending on their location and ecological attributes.

Third, we found stronger spatiotemporal overlap among related mice. Indeed, kinship is an important organizing principle for many animal societies, with individuals preferentially associating with genetic relatives (Silk 2002). Non-random spatial associations among related individuals have also been reported for Alabama beach mice (*P. p. ammobates*) using trap data (Tenaglia et al. 2007). In Santa Rosa beach mice (*P. p. leucocephalus*), we find that related mice tend to sleep in the same burrow during the day and use overlapping burrow networks at night. One limitation of our study is that we cannot determine which mouse dug which burrow and therefore cannot directly compare the costs of burrow construction to the benefits of burrow use. Because digging is energetically costly (Reichman and Smith 1990; Vleck 1979), it raises the question of why mice might build multiple burrows. One possibility is that beach mice invest in digging and maintaining multiple burrows throughout their home range because they receive inclusive fitness benefits when those burrows are also used by kin. Longer-term monitoring of burrow-use patterns, coupled with measurements of survival and reproduction, are necessary to fully address this question.

Here, by employing continuous recordings of beach mouse burrow use in the wild, we suggest that selection for long burrows may be driven, at least in part, by thermal buffering in their exposed habitat, which may be especially important for pup survival. Moreover, by combining these data with genetic analyses, we demonstrate that genetically related beach mice use overlapping networks of spatially clustered burrows, underscoring the centrality of burrows in the social life of these monogamous mice. Overall, our findings highlight how developing new automated methods to non-invasively monitor animals in the wild can provide novel insights into their otherwise secretive behavior.

## Acknowledgements

We thank Erica Laine and Glenn Barndollar of Eglin Air Force Base for logistical support and project advice as well as Jeff Gore of Florida Fish and Wildlife for project advice. We thank Caitlin Lewarch for early help in the field; Ed Soucy of the Harvard Center for Brain Science Neuroengineering Core for help designing and assembling RFID readers; Mark Omura, Judy Chupasko, and Breda Zimkus for assistance with specimen accessions; Paul Moorcroft, Nathan Ranc, and members of the Hoekstra Lab for helpful discussions. This work was supported by a Grant-in-Aid of Research from the American Society of Mammalogists and a Natural Science and Engineering Research Council of Canada Postgraduate Scholarship to N.L.B. H.E.H. is an Investigator of the Howard Hughes Medical Institute.

## Author Contributions

N.L.B. and H.E.H. designed the experiments. N.L.B., J.T.G., C.K.H., T.B.W. and N.A.S. conducted the field experiments. J.M.L. designed the targeted genotyping assay. N.L.B. analyzed the data. N.L.B. and H.E.H. wrote the manuscript with input from all authors.

## Materials & Correspondence

Correspondence and material requests should be addressed to H.E.H.

## Data Availability

All data and code available at https://github.com/nicolebedford/BeachMouse_RFID

