## Supplementary Material for "Automated tracking reveals the social network of beach mice and their burrows"

### November 2017 Field Season: Extended Methods

#### <u>Extended Methods</u>

All procedures were approved by the IACUC at Harvard University (Protocol No. 27-17), the Florida Fish and Wildlife Conservation Commission (Permit No. LSSC-15-00094), and the Eglin Air Force Base RC3 Committee.

*<u>Phase I: Capture-recapture (November 2-17, 2017)</u>*

In November 2017, we established a 70 x 9 grid of 630 Sherman folding traps (H.B. Sherman Traps, Inc., Tallahassee, FL) in the frontal dunes of Santa Rosa Island, Florida (30°23’42” N, 86°43’27” W) (Fig. S1A, B). Traps were spaced 10 m apart in both directions. The final trap grid measured 690 x 80 m (55,200 $m^2$) and ran parallel to the Gulf of Mexico. We set traps an hour before sunset each night (approx. 5 PM) and checked an hour after sunrise (approx. 6 AM) the next morning. Traps were baited with Pennington Select Black Oil Sunflower Seed (Pennington Seed Inc., Madison, GA).

Upon capture, we determined the sex, age, and reproductive status of all trapped mice. We measured body mass using a Pesola® LightLine Spring Scale (Forestry Suppliers, Inc., Jackson, MS). We measured total length, tail length, ear length, and hindfoot length using a ruler. Next, we briefly anesthetised mice with isoflurane and injected an 8 mm 125 kHz RFID tag (Freevision Technologies Co., Shanghai, China) under the scruff (i.e., between the shoulder blades). We also collected a small clip of ear tissue for subsequent DNA extraction. Finally, we released each mouse at its original trap location and noted if the animal entered a nearby burrow upon release. In this way, we identified putative beach mouse burrows for RFID monitoring.

We trapped for 16 consecutive nights (November 2-17, 2017) and caught 36 mice (see Table S1 for details). Our capture-recapture results are summarized in Fig. S1D. After 16 nights of grid trapping, we set 3-4 traps near the entrances of putative beach mouse burrows for two additional nights (November 18-19, 2017). With this targeted approach we trapped 7 new animals: 1 adult and 6 juveniles, 2 of which were too small to tag. We did not include these captures in our spatially explicit capture-recapture (SECR) models. In total, we trapped 43 mice, 39 of which were successfully tagged and released.

**Table S1.** November 2017 capture-recapture summary.

| **Night** | **1** | **2** | **3** | **4** | **5** | **6** | **7** | **8** | **9** | **10** | **11** | **12** | **13** | **14** | **15** | **16** | **Total** |
|---|---|---|---|---|---|---|---|---|---|---|---|---|---|---|---|---|---|
| Traps set | 80 | 328 | 414 | 414 | 414 | 495 | 495 | 495 | 495 | 495 | 495 | 495 | 495 | 495 | 243 | 135 | 6483 |
| Captures | 4 | 4 | 7 | 8 | 3 | 7 | 7 | 6 | 6 | 10 | 10 | 14 | 15 | 16 | 6 | 5 | 128 |
| New | 4 | 2 | 2 | 3 | 0 | 1 | 2 | 4 | 3 | 4 | 1 | 3 | 4 | 1 | 2 | 0 | 36 |
| Losses | 1 | 0 | 0 | 1 | 0 | 0 | 0 | 0 | 0 | 0 | 0 | 0 | 0 | 0 | 0 | 0 | 2 |
| Cumulative | 4 | 6 | 8 | 11 | 11 | 12 | 14 | 18 | 21 | 25 | 26 | 29 | 33 | 34 | 36 | 36 | 36 |

*<u>Phase II: RFID monitoring (November 20-30, 2017)</u>*

We identified burrows for RFID monitoring by noting which burrows mice entered upon release. Ghost crabs (*Ocypode quadrata*) are abundant in the foreshore and excavate burrows that appear superficially similar to those of beach mice (Hill and Hunter 1973). We found that mice often entered ghost crab burrows upon release, but later emerged and darted into a different burrow. Active beach mouse burrows were distinguished from ghost crab burrows (which are likely used as temporary refugia) by trial and error. Specifically, we installed custom-built RFID readers at putative beach mouse burrow entrances and kept readers in place for a minimum of 2 nights. If no activity was detected after 2 consecutive nights, the RFID reader was moved to a new burrow. Over 11 consecutive nights, we recorded RFID activity at 40 separate burrows, but not all burrows were active on all nights (see Table S2 for details). Out of 39 mice that were successfully tagged and released in Phase I, we detected 32 at our RFID-monitored burrows in Phase II.

**Table S2.** November 2017 RFID summary.

| Night | 1 | 2 | 3 | 4 | 5 | 6 | 7 | 8 | 9 | 10 | 11 | Total |
|---|---|---|---|---|---|---|---|---|---|---|---|---|
| Active RFID readers | 26 | 24 | 27 | 27 | 27 | 27 | 28 | 28 | 23 | 22 | 22 | 40 |
| No. mice detected | 20 | 21 | 19 | 20 | 23 | 21 | 20 | 18 | 21 | 20 | 17 | 32 |
| Total activity (no. reads) | 1152 | 1003 | 798 | 783 | 1137 | 876 | 621 | 382 | 676 | 545 | 338 | 8311 |

*Custom RFID readers*

We designed and built custom, low-cost RFID readers to track mouse activity at burrow entrances. We used a Teensy 2.0 USB development board (PJRC Electronic Projects) to run a custom Arduino script (version 1.6.1). Our readers consisted of a 125 Khz RFID module (Seeed Technology Co., Shenzhen, China), a ChronoDot V2.1 Ultra-precise Real Time Clock (Macetech LLC, Vancouver WA), and a microSD breakout board (Adafruit Industries, LLC, New York NY). All components were soldered to a custom PCB board (ExpressPCB LLC, Mulino OR) and powered by a 7 Ah 12 Volt Battery (Universal Power Group, Inc., Coppell TX). We built custom antennas by wrapping copper magnet wire (Belden Inc., St. Louis MO) around a laser-cut acrylic frame (5 cm diameter) and installing galvanized bolts to hold the antenna in place at the burrow entrance (Fig. 1B). We waterproofed the antenna with Permatex® Clear RTV Silicone Adhesive Sealant (Illinois Tool Works Inc., Glenview IL) and stored the RFID reader and battery in a waterproof case (Plano Molding Co., Plano IL). In total, we assembled 30 custom RFID readers.

*Home range size and population density*

To estimate home range size and population density, we ran a spatially explicit capture-recapture (SECR) analysis using the *secr* package in R (Efford 2018). We included in our model all capture events from nights 3-14 when more than 400 traps were set. We used the “suggest.buffer” function to determine a suitable habitat mask (80 m) around the periphery of the 690 x 80 m trap grid (Fig. S1C). We then fit a half-normal spatial detection model by maximum likelihood using the “secr.fit” function, which estimates two parameters: $g_0$ and $\sigma$ (Table S3). The capture probability at the home range center is denoted by the intercept ($g_0$), and the scale parameter ($\sigma$) describes how this probability decays with increasing distance from the home range center (Fig. S1E). We estimated the 95% home range radius from $\sigma$ using the “circular.r” function.

**Table S3.** November 2017 SECR model results.

| Value | estimate | s.e. |
|---|---|---|
| Prob. of capture at activity center ($g_0$) | 0.019 | 0.003 |
| Scale parameter ($\sigma$) | 29.97 | 2.46 |
| 95% home range radius (m) | 67.26 | - |
| Home range size ($m^2$) | 14,212 | - |
| Density (mice/$km^2$) | 254.0 | 48.0 |

*Burrow attributes*

At the end of the study, we measured various physical and ecological attributes of the RFID-monitored burrows (Fig. S2). To measure the length of the entrance tunnel, we inserted flexible plastic tubing into the burrow and recorded the maximum distance. We took the average of four inclinometer readings to determine the mean slope of the dune at the burrow entrance and took the average of three compass readings to determine the heading of the burrow entrance. We also recorded the elevation (in feet above sea level) of each burrow entrance. Finally, to determine the extent of vegetative cover, we photographed the burrow entrance centered inside a 50 x 50 cm quadrat. A researcher blind to burrow identity quantified percent cover in FIJI (Schindelin et al. 2012).

*Burrow temperature profiles*

We recorded daily temperature fluctuations using HOBO® U23 Pro v2 data loggers (Onset Computer Co., Bourne, MA). In November 2015, we constructed artificial burrows by coring sand plugs of various lengths (e.g., 10, 40, 60, and 80cm) from the sides of dunes at a 25-degree downward angle. We constructed and recorded from artificial burrows at 7 separate dune sites from November 2-17, 2015 (Fig. S3A). At each site, we placed one HOBO® data logger at the dune surface to record local ambient temperature. From May 4-17, 2016, we recorded from natural burrows located outside the trap grid at 2 separate dune sites (Fig. S3C). From November 4-29, 2017, we recorded from natural burrows located outside the trap grid at 3 separate dune sites (Fig. S3D). At the end of each recording session, we measured the length of the entrance tunnel by inserting flexible plastic tubing into the burrow and recording the maximum distance.

To determine a standard metric of temperature fluctuations across burrows measured on different dates in different seasons, we first detrended the data by regressing temperature against time. We then extracted the amplitude envelope of the residual temperature fluctuations using the Hilbert transform (*seewave* package in R). We then tested for an effect of burrow length on the mean value of the amplitude envelope using linear regression (Fig. 1G).

*RFID read filtering*

We removed trains of consecutive RFID reads occurring at 1Hz that might indicate an animal pausing at the burrow entrance. For all 1 Hz trains ≥ 3 reads, we kept the first and last read of the train and removed all intervening reads. In all, we removed 794 reads for a final set of 7517 RFID reads over 11 nights. In November 2017, we detected RFID activity for 32 mice at 40 different burrows.

*Hierarchical clustering*

To identify burrows with similar visitation profiles, we first determined the fraction of each burrow's total RFID activity comprised by each mouse. We then scaled and centered these data and calculated a dissimilarity matrix based on Euclidean distance. Next, we used Ward's minimum variance method to perform agglomerative hierarchical clustering.

*Spatiotemporal overlap*

To determine the extent of spatiotemporal overlap between mice, we developed an overlap index (OI). We defined $OI_{i,j}$ as the percentage of total RFID activity for mouse *i* that occurs at the same burrow and within the same 90-min time bin as mouse *j*. OI is not symmetrical (i.e., $OI_{1,2} \neq OI_{2,1}$) because it is expressed as a percentage of the focal mouse's (mouse *i*) total activity. We used the "SlidingWindow" function from the *evobiR* package in R (Blackmon and Adams 2015) to calculate OIs for a range of time bins, from 15 min to 14 hrs, using a step size of ½ the time bin size. For each time bin, we then calculated the network load (Psorakis et al. 2015), defined as the number of observed links in the network divided by the total number of possible links (i.e., the fraction of all possible pair combinations with non-zero OIs). We selected an appropriate time bin for determining spatiotemporal overlap by plotting network load as a function of time bin size. We approximated 1000 points along this curve using the function "approx" in R and identified the elbow of the curve (89 min) by calculating the maximum second derivative. We ran all downstream analyses on OIs calculated with a 90-min sliding window time bin.

*Genotyping and genetic relatedness*

To identify polymorphic markers for *P. p. leucocephalus*, we re-analyzed the gDNA data of representative individuals from a previous study (Domingues et al. 2012) (Short Read Archive accession number SRP010898). This dataset comprised data from 20 individuals sampled at ~5000 1.5kb targeted non-coding regions randomly distributed across the genome (see Domingues et al. 2012 for capture array design details). We aligned raw gDNA paired-end reads to an *in-house* assembly of the *Peromyscus polionotus* genome (Genbank assembly accession: GCA_003704135.1) using BWA (version 0.7.12) (Li and Durbin 2009). The resulting alignment files were pre-processed according to GATK Best Practices and used to perform SNP and INDEL discovery and genotyping across all 20 samples separately using HaplotypeCaller as implemented in GATK version 3.7 (McKenna et al. 2010; Van der Auwera et al. 2013). Default parameters were used, with the exception of the prior for heterozygosity, which was set to 0.005. We then performed joint genotyping on the gVCF files produced by HaplotypeCaller and filtered the resulting variant call set using BCFtools version 1.7 (Li et al. 2009) with standard hard filtering parameters (SNP: QD < 2.0, FS > 60.0, MQ < 40.0, SOR >3.0, MQRankSum < -12.5, ReadPosRankSum < -8.0; INDEL: QD < 2.0, FS > 200.0, ReadPosRankSum < -20.0). In addition, we filtered out variants called at GQ < 20 with a DP < 7, excluded sites with missing genotypes in > 10 individuals, and selected only biallelic variant sites. The final set of polymorphic sites were identified using the following criteria:

1. A site is considered variable if at least one sample of each category (hom_ref, hom_alt, het) is present. In addition, not all samples should be homozygous for either reference or alternate allele (1 < AC < 35).

2. Sites should not have an abnormally high number of heterozygous genotypes, as an excessive number of heterozygotes may reflect technical errors. We therefore excluded variants with ExcessHet ≥ 7.
3. To identify the most variable sites, we calculated a "variability score" defined as the product of the number of samples represented in each genotype category. For a given site, the higher the score, the more mixed the genotype profile is.
4. To avoid variants that were on the same segment, we excluded variants that were less than 200bp away from the nearest neighbour.

The resulting set of variants passing these filters comprised 294 sites. We extracted from the genome nucleotide sequences 100bp upstream and downstream of the 140 most variable sites (ranked according to our custom metric). We excluded markers that had more than one hit in the genome, as revealed by a local search using BLAST as implemented in Biopython (Cock et al. 2009). We selected a final set of 110 loci for designing genotyping probes.

We designed PCR primers for these target regions using BatchPrimer3 v1.0 (https://probes.pw.usda.gov/batchprimer3/). We used the default settings for general primers and selected final amplicon sizes between 80 and 90bp. We extracted DNA from ear tissue samples using an AutoGenprep 965 (Autogen, Holliston MA). Multiplex amplification of the targets was done by Floodlight Genomics, LLC (Knoxville, TN) using an optimized Hi-Plex approach as part of a no-cost Educational and Research Outreach Program (Nguyen-Dumont et al. 2013). Pooled barcoded amplicons were sequenced on an Illumina HiSeq X Five in a 2 x 150bp paired-end run. The sample-specific sequences were aligned to the target sequences used for primer design with CLC Genomics Workbench (Version 9.5.3). We assigned genotypes for loci with >10X coverage and used an alternate allele frequency cut-off of 15% to assign heterozygous calls. Of the 110 sequenced loci, 17 were monomorphic. With the remaining 93 polymorphic sites, we estimated pairwise kinship coefficients using "--relatedness2" in VCFtools (Manichaikul et al. 2010). We multiplied the kinship coefficient by 2 to obtain genetic relatedness (*r*).

<u>Social network analysis</u>

We constructed network graphs using the *igraph* package in R (Csardi and Nepusz 2006). For our network graphs, we used a Fruchterman-Reingold force-directed layout algorithm, which places nodes (i.e., mice) that share more connections closer together. We first generated a network graph of kinship coefficients, depicting genetic relationships between mice. We then saved this force-directed layout and applied it to a network graph of OIs to facilitate comparison between networks. For both networks, we calculated degree centrality (i.e., the number of direct connections) for each node using "centr_degree" in *igraph*.

<u>Daytime burrow analysis</u>

To determine where mice slept during the day, we extracted the first and last RFID read of each mouse's nocturnal activity period. We restricted our analysis to a subset of 12 mice that were detected on all 11 nights of the RFID study. We also removed "first exit" reads that occurred after 12 AM and "last entry" reads that occurred before 12 AM. For each mouse, we asked if the first read at dusk occurred at the same burrow as the last read the previous dawn. If so, the mouse was assigned a daytime burrow for that day. If not, the daytime burrow was not determined. On average, mice emerged from their daytime burrow at 5:48 PM and returned to their daytime burrow at 5:14 AM (Fig. S4B). At the midpoint of the study, sunset and sunrise occurred at 4:46 PM and 6:20 AM CST, respectively.

*Statistics*

All data analyses and statistics were performed in R (version 3.5.0). We ran linear regression models to test for effects of percent cover (Fig. 1E) and entrance tunnel length on nightly RFID activity. To obtain nightly RFID activity for each burrow, we averaged the total number of RFID reads across all nights for which the burrow was active. We then $\log_2$ transformed this value to improve normality. We also used linear regression to test for an effect of entrance tunnel length on the percentage of nightly RFID activity comprised by juvenile *versus* adult mice (Fig. 1F) and for an effect of entrance tunnel length on temperature fluctuation in natural burrows (Fig. 1G). We used an ANOVA, followed by Tukey's test to ask if artificial burrows of various lengths differed in their capacity for thermal buffering (Fig. S3B).

We used linear regression to test for an effect of kinship coefficient on overlap index, or OI (Fig. 4C). Additionally, we used a Komolgrov-Smirnov (K-S) test to ask if OI distributions based on real and randomized data differed significantly. We used another K-S test to ask if kinship coefficient distributions differed for mice sleeping in shared *versus* separate daytime burrows. Finally, we used simple linear models to test for differences in degree centrality (derived from network graphs for genetic relatedness and OI) between juveniles and adults and between males and females.

### May 2016 Field Season: Methods and Results

#### Methods

*Phase I: Capture-recapture (April 30 - May 10, 2016)*

In April and May 2016, we established a 60 x 7 grid of 420 Sherman folding traps (H.B. Sherman Traps, Inc., Tallahassee, FL) in the frontal dunes of Santa Rosa Island, Florida. To minimize "edge effect" we established a trap grid approximately 16 times larger than the average home-range size (Bondrup-Nielsen 1983). Traps were spaced 15 m apart in both directions. The final trap grid measured 885 x 90 m (79,650 $m^2$) and ran parallel to the Gulf of Mexico. We set traps an hour before sunset each night and checked at sunrise the next morning. Traps were baited with Pennington Select Black Oil Sunflower Seed (Pennington Seed Inc., Madison, GA). Mice were processed as described for the November 2017 field season.

**Table S4.** May 2016 capture-recapture summary.

| **Night** | **1** | **2** | **3** | **4** | **5** | **6** | **7** | **8** | **9** | **10** | **Total** |
|---|---|---|---|---|---|---|---|---|---|---|---|
| Traps set | 210 | 420 | 420 | 420 | 420 | 420 | 420 | 420 | 420 | 420 | 3990 |
| Captures | 3 | 6 | 2 | 7 | 12 | 12 | 16 | 14 | 16 | 24 | 112 |
| New | 3 | 5 | 2 | 3 | 5 | 1 | 3 | 4 | 6 | 8 | 40 |
| Losses | 1 | 0 | 0 | 0 | 0 | 0 | 0 | 0 | 0 | 0 | 1 |
| Cumulative | 3 | 8 | 10 | 13 | 18 | 19 | 22 | 26 | 32 | 40 | 40 |

*Phase II: RFID monitoring (May 11-18, 2016)*

Over 7 consecutive nights, we recorded RFID activity at 29 separate burrows, but not all burrows were active on all nights (see Table S5 for details). Out of 39 mice that were successfully tagged and released in Phase I, we detected 31 at our RFID-monitored burrows in Phase II.

**Table S5.** May 2016 RFID summary.

| Night | 1 | 2 | 3 | 4 | 5 | 6 | 7 | Total |
|---|---|---|---|---|---|---|---|---|
| Active RFID readers | 10 | 15 | 15 | 15 | 20 | 16 | 17 | 29 |
| No. mice detected | 16 | 19 | 18 | 18 | 22 | 19 | 21 | 31 |
| Total activity (no. reads) | 190 | 599 | 929 | 222 | 364 | 489 | 237 | 3030 |

*SECR model*

To estimate home-range size and population density, we ran a spatially explicit capture-recapture (SECR) analysis using the *secr* package in R. We included in our model all captures events from 10 consecutive trap nights. We used the “suggest.buffer” function to determine a suitable habitat mask (116 m) around the periphery of the 885 x 90 m trap grid. We then fit a half-normal spatial detection model by maximum likelihood using the “secr.fit” function, which estimates two parameters, $g_0$ and $\sigma$ (Table S6). The capture probability at the home range center is denoted by the intercept ($g_0$), and the scale parameter ($\sigma$) describes how this probability decays with increasing distance from the home range center. We estimated the 95% home range radius from $\sigma$ using the “circular.r” function.

**Table S6.** May 2016 SECR model results.

| Value | estimate | s.e. |
|---|---|---|
| Prob. of capture at activity center ($g_0$) | 0.029 | 0.005 |
| Scale parameter ($\sigma$) | 44.53 | 2.95 |
| 95% home-range radius (m) | 99.94 | - |
| Home-range size ($m^2$) | 31,375 | - |
| Density (mice/$km^2$) | 157.6 | 26.9 |

*RFID read filtering*

We removed trains of consecutive RFID reads occurring at 1Hz that might indicate an animal pausing at the burrow entrance. For all 1Hz trains ≥ 3 reads, we kept the first and last read of the train and removed all intervening reads. In all, we removed 249 reads for a final set of 2781 RFID reads over 7 nights. In May 2016, we detected RFID activity for 31 mice at 29 different burrows.

### Results

*Population sampling*

Using a spatially explicit capture-recapture model, we estimated a population density of 158 ± 27 mice/$km^2$. The probability of trapping a mouse at its home-range center was 2.9 ± 0.5% and the mean home-range size estimate was 31,735 $m^2$. This estimate is considerably larger than our home-range size estimate of 14,212 $m^2$ for November 2017, which may reflect seasonal differences in movement patterns.

*Patterns of burrow use*

As in November 2017, we found that mice used multiple burrows, and that burrows were used by multiple mice. Specifically, a single mouse would visit between one and four burrows per night, and a single burrow would be visited by one to four mice per night. Likewise, over the course of the 7-night sampling period, individual mice visited between one and seven burrows (median = 3 burrows) and individual burrows were visited by one to seven mice (median = 3 mice).

*Spatiotemporal overlap*

We found evidence of spatiotemporal overlap, which we defined as non-zero OI values, for 39/465 possible pairs (8%). We identified four groups of two and three groups of three mice that showed strong spatiotemporal associations (white boxes, Fig. S5A). The maximum OI was 3.33% and the mean OI (excluding zeros) was 0.80%. By contrast, when we shuffled the data, we found evidence of spatiotemporal overlap for 100% (465/465) of pairs in the randomized dataset. However, the magnitude of overlap was much larger for real compared to randomized data (Fig. S5B). In the randomized data, the maximum OI was 0.95% and the mean OI (excluding zeros) was 0.42%. Consistent with our conclusions for November 2017, these findings suggest that when mice do overlap, they show stronger spatiotemporal associations than expected by chance.

*Genetic relatedness*

We made high-confidence SNP calls for 27/31 mice and estimated coefficients of relatedness ($r$) for all possible pairs of mice ($n$ = 351). Only 35/351 pairs showed spatiotemporal overlap at RFID-monitored burrows. Of these 35 overlapping mice, 26 had $r > 0$ and 9 were unrelated ($r = 0$). Thus, given that two mice overlapped, the probability that they were genetic relatives ($r > 0$) was 74% (Binomial test, $P$ = 0.006, 95% $CI$ = 0.57, 0.88).

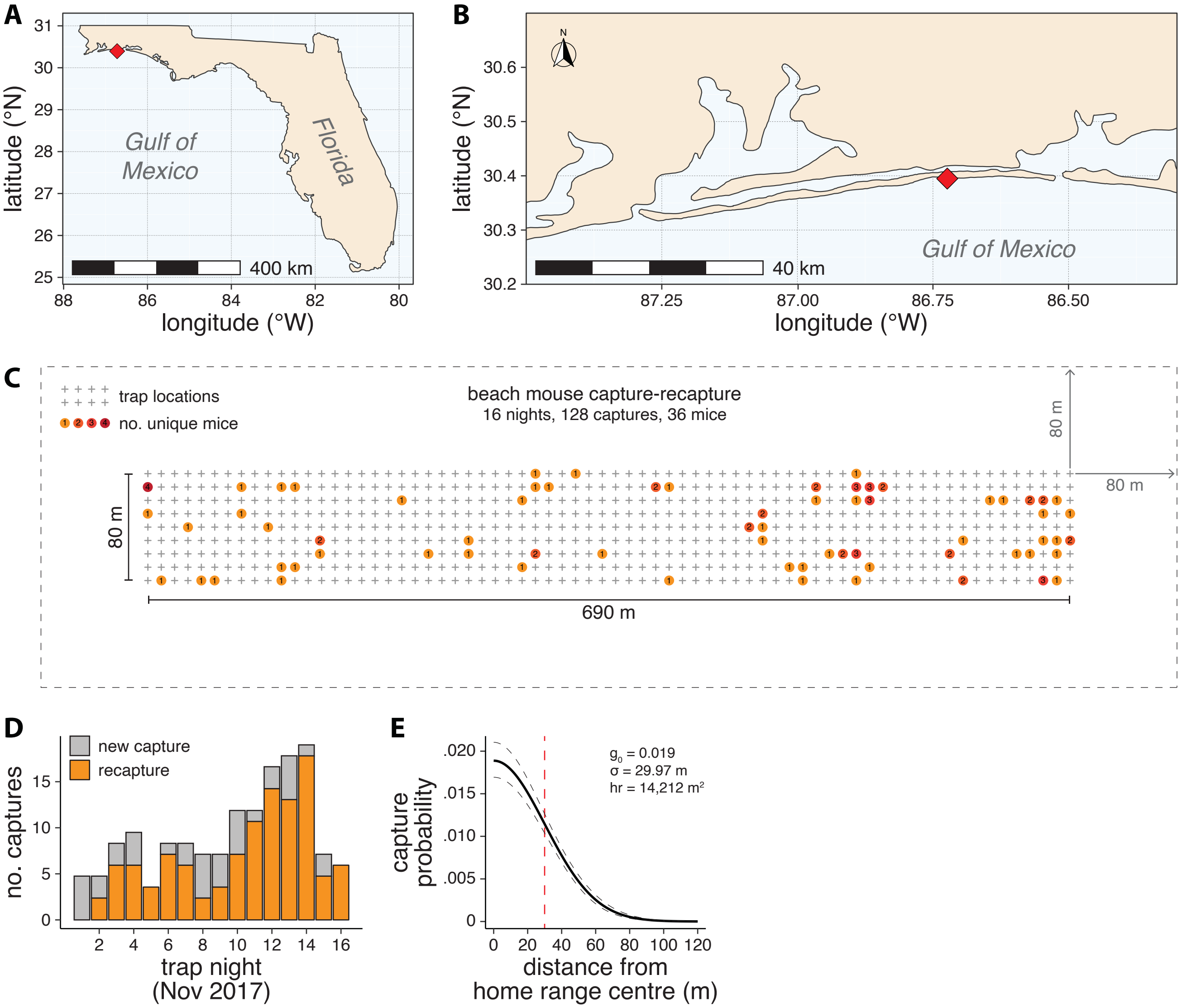


**Figure S1: Capture-recapture study details and model results. (A)** Map of Florida, with field site location indicated (red diamond). **(B)** Map of Santa Rosa Island, with field site location indicated (red diamond). **(C)** Schematic of 690 x 80 m trap grid, with 630 traps (grey crosses) spaced 10 m apart in both directions. Colored circles indicate the number of unique mice caught at each trap location. Dashed grey line denotes the habitat mask boundary (80 m in both directions). **(D)** Capture-recapture data for 16 consecutive trap nights (November 2-17, 2017). **(E)** Spatially explicit capture-recapture (SECR) model results. Half-normal detection function describing capture probability as a function of distance from the home range centre. Dashed grey line represents the standard error of the model estimate, and the dashed red line indicates the scale parameter value (σ). g0 = intercept, σ = scale parameter, hr = 95% home range radius.

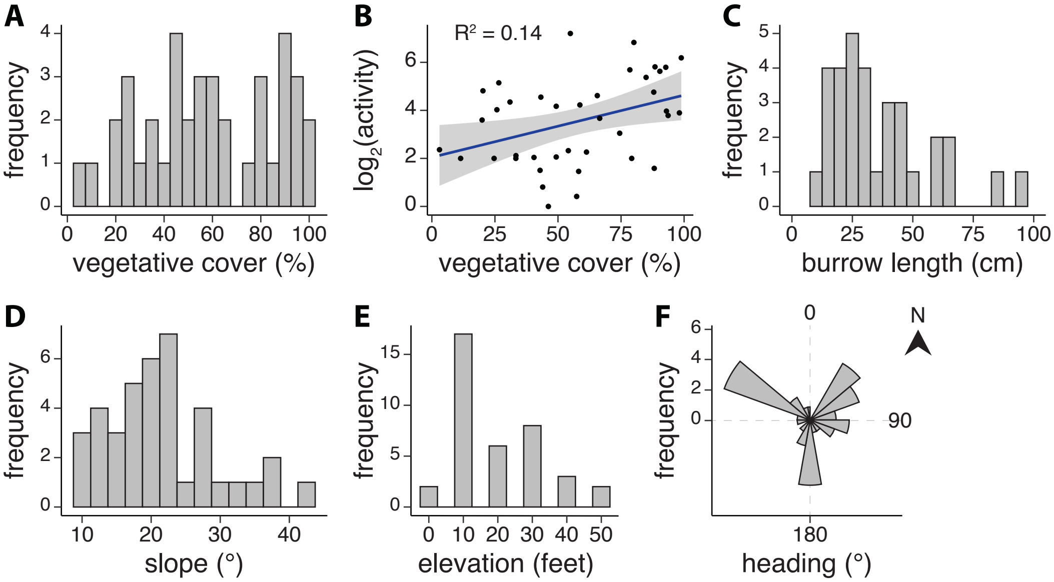


**Figure S2: Ecological and physical attributes of RFID-monitored burrows. (A)** Burrow site vegetative cover (mean = 58%). **(B)** Relationship between percent cover and average nightly RFID activity. **(C)** Burrow entrance tunnel length (mean = 37 cm). **(D)** Burrow site slope (mean = 22°). **(E)** Burrow site elevation (mean = 20 feet above sea level). **(F)** Burrow entrance headings (mean = 178°). Zero degrees indicates true North.

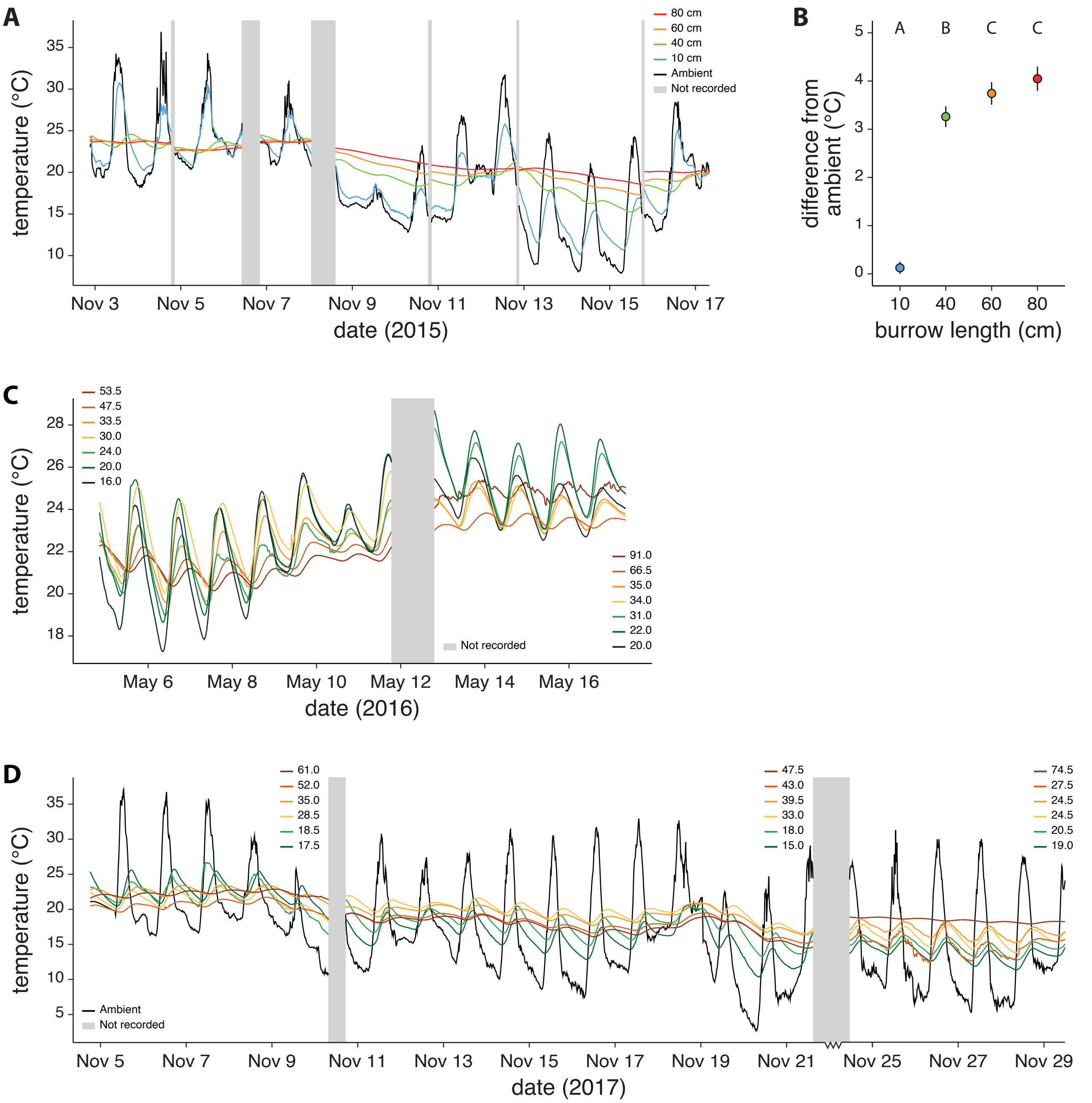


**Figure S3: Temperature profiles of artificial and natural burrows. (A)** Temperature profiles of artificial burrows at 7 separate dune sites in November 2015. **(B)** Average difference from ambient temperature for all 10, 40, 60 and 80 cm burrows. Groups with different letters are significantly different (Tukey's test). **(C)** Temperature profiles of natural burrows of varying lengths at 2 separate dune sites in May 2016. **(D)** Temperature profiles of natural burrows of varying lengths at 3 separate dune sites in November 2017. Grey bars denote times when data loggers were moved between sites. Ambient temperature is shown in black.

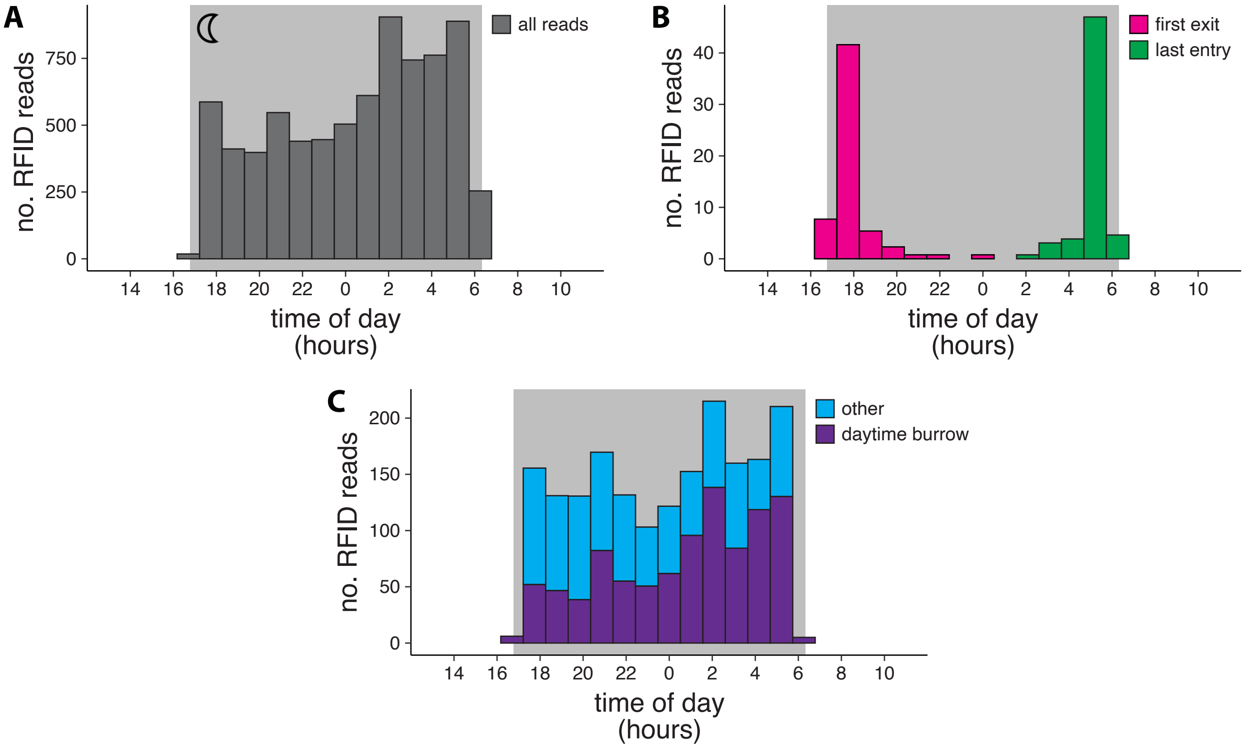


**Figure S4: Timestamp distributions of RFID reads. (A)** Timestamps of all RFID reads over the 11-night sampling period. **(B)** Timestamps of the first burrow exit and last burrow entry of the night, per mouse. **(C)** Timestamps of RFID reads recorded at the daytime burrow (i.e. the burrow where the mouse slept) *versus* other burrows in the home range. RFID reads from a subset of 12 mice for whom daytime burrow location was determined are included in **(B)** and **(C)**. For all plots, the shaded grey area denotes nighttime. The sun set at 16:46 and rose at 06:20 CST at the midpoint of our RFID-monitoring period (November 20-30, 2017).

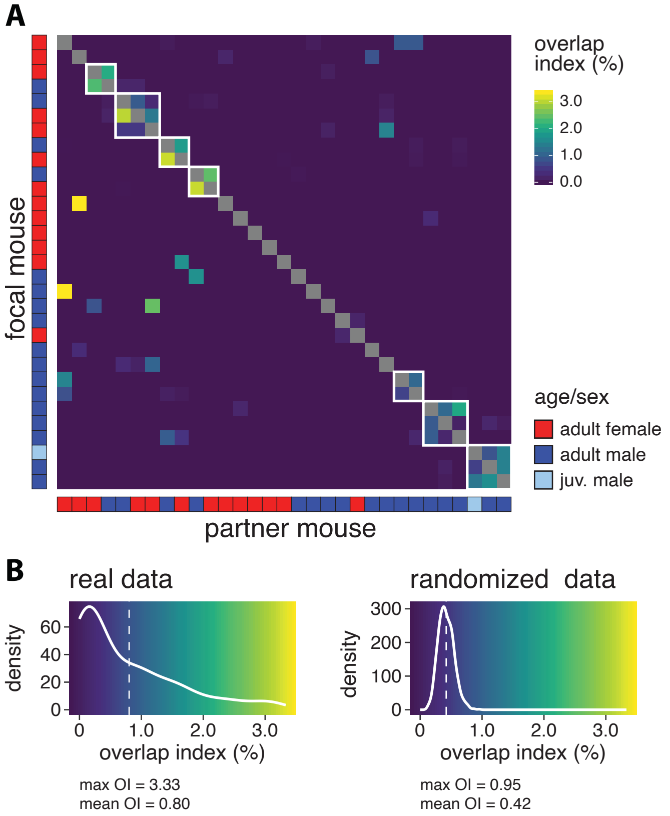


**Figure S5: Mice show stronger spatiotemporal overlap than expected by chance. (A)** Overlap index (OI) matrix for all possible focal and partner mouse pairs. For each focal mouse, OI indicates the percentage of total RFID activity that occurs at the same burrow and within the same 90-min time bin as the partner mouse. **(B)** Distributions of non-zero OIs from real data (left) and randomized data (right). Dashed white lines indicate distribution means. Data from May 2016 are shown.
